# Lonely plants in arid land are functionally hyperdiverse

**DOI:** 10.1101/2025.02.21.639323

**Authors:** Pierre Liancourt, Raphael Martin, Yoann Le Bagousse-Pinguet, Fernando T. Maestre, Miguel Berdugo, Manuel Delgado-Baquerizo, David J Eldridge, Hugo Saiz, Santiago Soliveres, Enrique Valencia, Nicolas Gross

## Abstract

In Gross et al.^1^ we produced the largest ever standardized dryland plant trait database including 133,769 trait measurements from 301 perennial plant species surveyed across 326 plots and six continents. Our findings indicate that arid and hyper-arid drylands act as a global reservoir of plant phenotypic diversity, challenging the common assumption that harsh environmental conditions reduce plant trait diversity. Tordoni et al.^2^ speculate that the larger phenotypic diversity in harsh environments found in our study is overestimated and misinterpreted. The re-analyses presented here further confirm that the patterns we originally reported are robust, and thus that the concerns from Tordoni et al. are not well-founded and do not apply to our study. We stand for the main conclusions of our study and maintain that lonely plants in arid land are functionally hyperdiverse.

## Main text

In Gross et al.^1^ we produced the largest ever standardized dryland plant trait database including 133,769 trait measurements from 301 perennial plant species surveyed across 326 plots and six continents. Our findings indicate that arid and hyper-arid drylands act as a global reservoir of plant phenotypic diversity, challenging the common assumption that harsh environmental conditions reduce plant trait diversity. Tordoni et al.^2^ speculate that the larger phenotypic diversity in harsh environments found in our study is overestimated and misinterpreted. They argue that (i) our sampling design is “unbalanced” because it overrepresents sites with aridity value > 0.7 (and therefore their trait variation and phenotypic diversity) and (ii) we should have consequently controlled for the number of species in our analyses. Strikingly, and despite the harsh criticisms, they conclude that our results remain detectable and clear, and that the findings we originally reported remain a challenge to conventional expectations, supporting our original conclusions. Although we do not deny the value of the argument, we show here that the concerns from Tordoni et al. are not well-founded and do not apply to our study.

We disagree with Tordoni et al. that our sampling is “unbalanced”. Our sampling scheme included more xeric (aridity value > 0.7) than mesic sites (aridity value < 0.7) because it reflects a biogeographical fact that we intended to capture in our design^3^ (see Fig. 1 in ref.^2^). This design mirrored the distribution of drylands observed along global aridity gradients^4^, and it is therefore a desirable feature of our survey. However, regarding the overrepresentation of xeric sites in our dataset, Tordoni et al. speculated that the greater number of sites sampled at the xeric end of the gradient led to an artificial inflation of the size of its species pool, which in turn resulted in the observed patterns. They used simulated data to illustrate the correlation between the number of species, the dimensionality of the hypervolume and its size for species pools ranging from 2 to 10 species (see Fig. 1 in Tordoni et al.). The relationship between species richness and functional diversity at a low number of species is well documented^5–7^. However, the number of species considered in each aridity window in our original study^1^ was far much higher (average number of species per window= 51; min= 32, max = 69, Extended Data Figure 1a). In this situation, the species richness-hypervolume relationship does not hold (Fig. 1a). These results clearly indicate that the criticisms raised by Tordoni et al. do not apply on the biogeographical scale used in our study (see red square in Fig. 1a).

**Figure 1.**
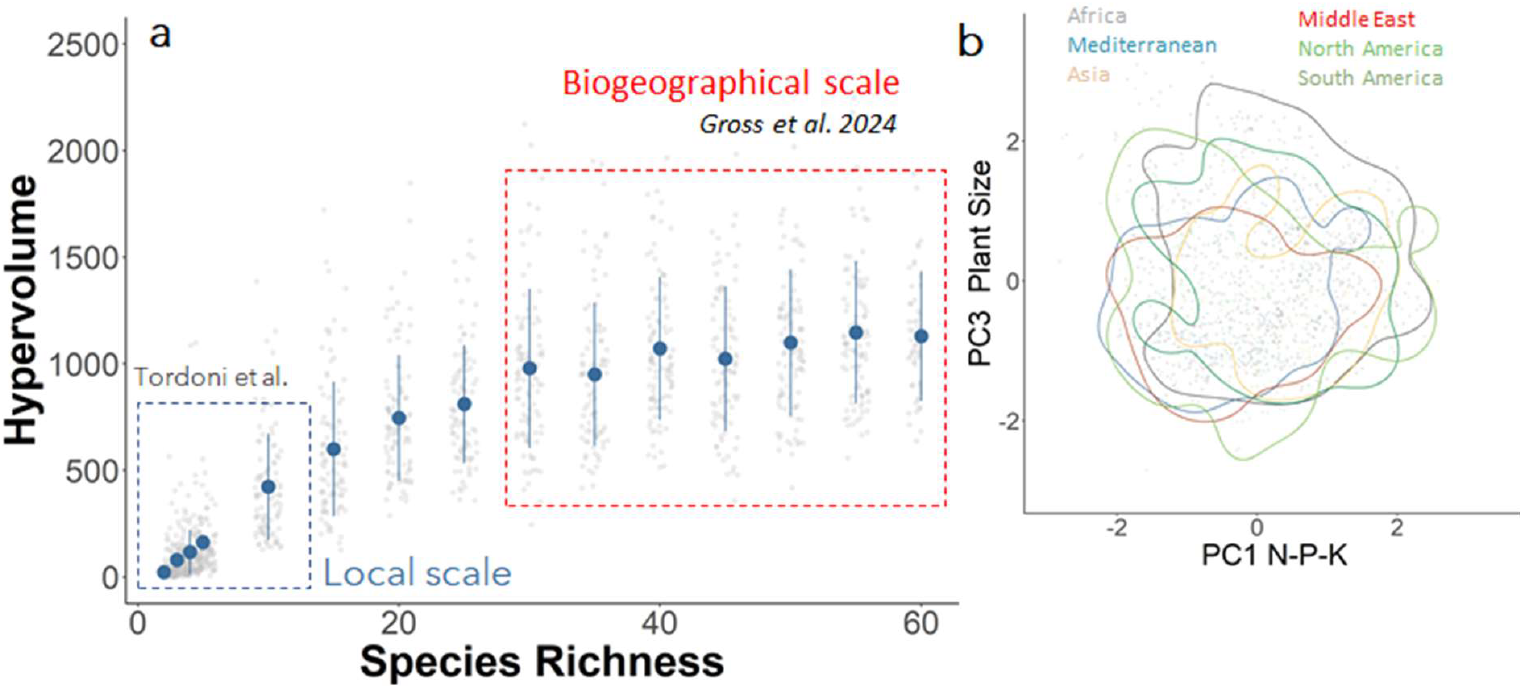
Species richness does not correlate with hypervolume at large biogeographical scales. (**a)** We generated the expected relationship between species richness and hypervolume by simulating plant assemblages ranging from 1 to 60 species. For each simulation, we randomly selected 100 individuals from the data of Gross et al. 2024 and computed hypervolumes using the code provided by the authors. A total of 100 simulations were run for each species richness level. Blue dots represented the mean hypervolume per species richness level, and the upper and lower lines show the standard error. Grey dots represent results for each simulation. We indicated the range of species richness considered in Figure 1 of Tordoni et at. (Tordoni et at., 2-10 species, blue dashed square) and the range of species richness observed in Gross et al. 2024 (Species richness > 30, red dashed square). Tordoni et at. report a positive relationship between species richness and hypervolume (see also Figure 1 in Tordoni et at.). This relationship is only observable at a low level of species richness, a number of species that can be observed at a local scale. At a large biogeographical scale such as that considered in Gross et al. 2024, this relationship is no longer observable. On average, our analysis indicates that sampling more than 30 species was sufficient to capture the size of the dryland plant hypervolume. Sampling more than 30 species does not increase the size of the hypervolume, indicating a strong functional overlap between dryland plant species. (**b)** Illustration of the functional overlap between flora from different biogeographical origins along the leaf carbon economy spectrum and the plant size spectrum (data extracted from Gross et al. 2024). Color lines show the 0.95 confidence interval for each biogeographical origin.

In essence, adding more species from different biogeographical origins did not influence our results because of the large functional overlap between these regions (Fig. 1b). Actually, when we fixed the number of species in the analyses rather than controlling for the number of individuals, as we did in our study^1^, we observed a stronger, rather than a weaker, increase in trait diversity in the most arid drylands evaluated (Fig. 2a). These analyses further indicate that such increase is not influenced by unbalanced sampling and species richness. Rather, they show - as discussed in our original study - that plant species occupied new regions of the trait space above an aridity threshold of 0.7, leading to higher dimensionality of the hypervolume and lower trait co-variation.

**Figure 2.**
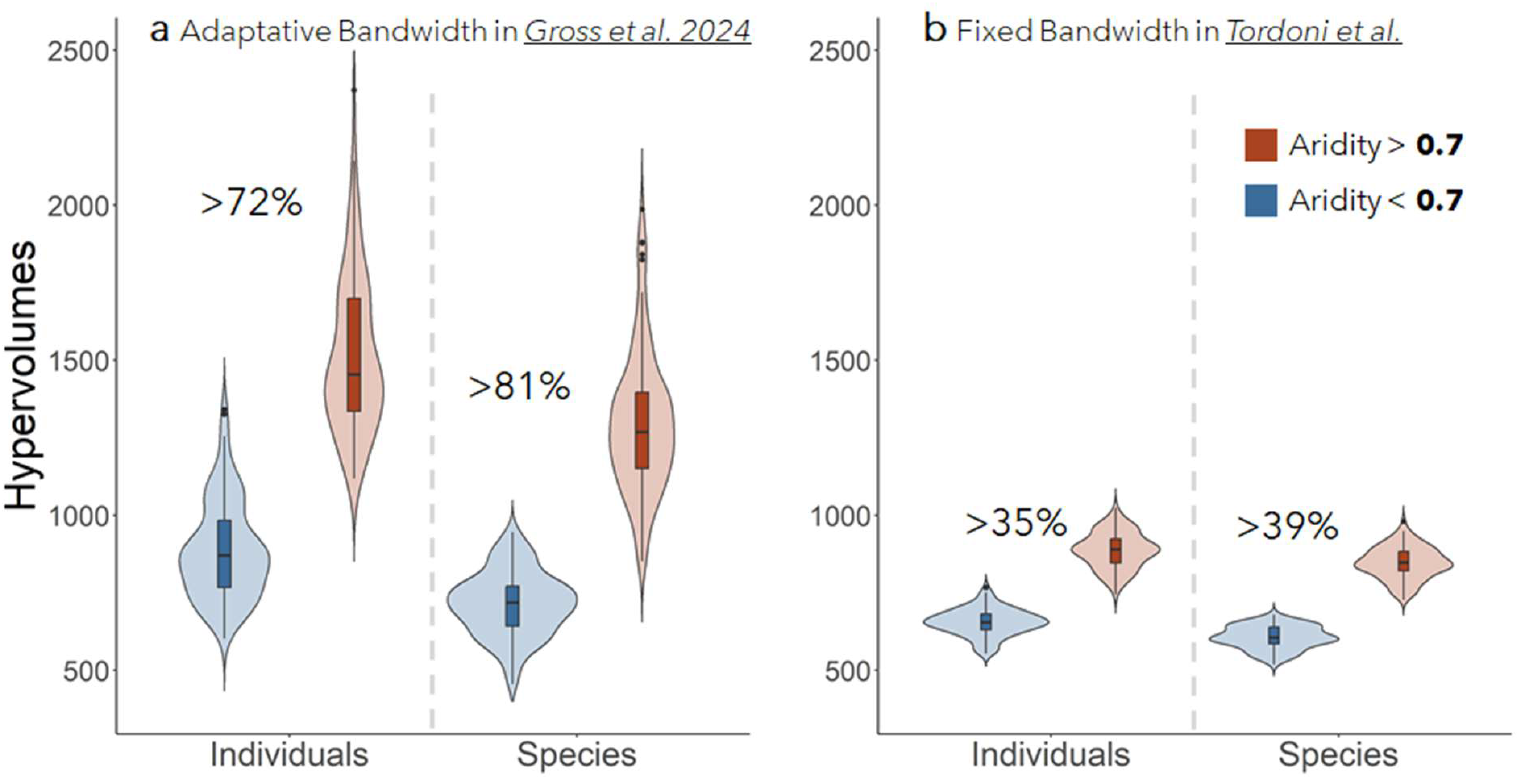
Hypervolume in areas above and below an aridity threshold of 0.7 computed with the adaptive bandwidth used in Gross et al. 2024 (**a**) and a fixed Bandwidth proposed by Tordoni et at. (**b**). We run the analyses with a fixed number of individuals or a fixed number of species (one individual per species). Hypervolumes with adaptive bandwidth were computed using the code provided by Gross et al. 2024. Hypervolumes with fixed bandwidth were computed using the code provided by Tordoni et al.. In all cases, bootstrapped values were generated using a random sampling of *n* = 80 observations for 100 times. Data are represented as boxplots where the middle line is the median, the lower and upper hinges correspond to the first and third quartiles, the upper and lower lines show the 0.95 confidence intervals. Fixing the number of species did not change the results presented in Gross et al. as we observed a 81 % increase of trait diversity in more xeric sites above the 0.7 aridity threshold. These results indicated that the result of Gross et al. 2024 is not biased by the number of species. Rather, it is explained by the appearance of novel plant strategies that occupied new regions of the trait space once the 0.7 aridity threshold is crossed. In contrast, the analyses using fixed bandwidth did not capture the global increase of trait diversity. This is so because the fixed bandwidth does not account for increasing dimensionality in the trait space along the aridity gradient surveyed (see main text).

Tordoni et at. proposed the use of a fixed bandwidth to control for species richness based on the assumption that differences in species richness along the aridity gradient surveyed biased our results. We chose an adaptive, rather than fixed, bandwidth as recommended in ref.^8^. This method allows us to effectively account for changes in the distribution of individual species within the trait space influencing the standard deviation on each trait axis^9^. When running the analyses using their approach, we found a lower global increase in trait hypervolume (< 40%, Fig. 2b) even after controlling for a fixed number of species. This is so because the fixed-bandwidth approach is not able to account for changes in the dimensionality of the trait space^8,9^ along the aridity gradient. Adopting their alternative approach would, in fact, have obscured a key result of our study, i.e., the increase in hypervolume dimensionality through the decoupling of macro-nutrients in plants observed when crossing an aridity threshold of 0.7.

Importantly, Tordoni et al. missed a key finding from our study: it is not aridity, but plant isolation that drives high levels of trait diversity in arid environments. We found that aridity and grazing had a similar effect on trait diversity because both decreased plant cover. We suggested that declining plant cover in the most arid and grazed regions of our planet promotes a “plant loneliness syndrome”, where the scattered plants across the most arid rangeland landscapes exhibit a wide phenotypic variability. This syndrome has also been observed outside dryland areas, as demonstrated in a recent study on glacier extinction^10^, and we proposed that it may be driven by the release of competitive interactions^11^ as individual plants have more space to thrive by avoiding competitive interactions. Regardless of the mechanisms involved, it is interesting to note that our design implicitly considered changes in plant cover through increasing grazing pressure at each level of aridity, resulting in a balanced sampling of species richness at low and high plant cover (Extended Data Figure 1b). In this context, our study provides a global “quasi-experimental” test of the role of plant cover in driving dryland plant phenotypic diversity.

Anticipating biodiversity responses to ongoing global change represents an immense challenge. For instance, investigating how species richness and trait diversity co-vary across drylands is critically needed to quantify the degree of functional redundancy, a key parameter for ecosystem resilience to global change drivers^12,13^. Also, the extent to which intraspecific trait variability contributes to global pattern of trait diversity remains largely unknown^14^. These questions are relevant but clearly out of the scope of our original study, and thus must be addressed in future studies. The re-analyses presented here further confirm that the patterns we originally reported are robust, and thus that the concerns from Tordoni et al. are not well-founded and do not apply to our study. We stand for the main conclusions of our study and maintain that lonely plants in arid land are functionally hyperdiverse.

## Author Contributions

All Co-authors discussed the ideas developed in the manuscript. P.L. R.M., Y.L.B.P and N.G. design the statistical analyses. R.M. and N.G. run the analyses. R.M. wrote the reproducible R code. P.L. and N.G. wrote the first draft with inputs from all authors.

## Acknowledgements

N.G. was supported by CAP 20-25 (16-IDEX-0001)). FTM is supported by King Abdullah University of Science and Technology (KAUST) and by the KAUST Climate and Livability Initiative. Y.L.B-P. was also supported by a Marie Sklodowska-Curie Actions Individual Fellowship (MSCA-IF) within the European Program Horizon 2020 (DRYFUN Project 656035).

## Supplementary Information

is not available for this paper.

## Data availability statement

All the data used in this paper are from Gross et al. 2024 and available here: https://doi.org/10.1038/s41586-024-07731-3

## Code availability statement

The data and code needed to reproduce the analyses are available in Nextcloud https://nextcloud.inrae.fr/s/y7kSyE69RWP22tD

---

**Extended Data Figure 1.**
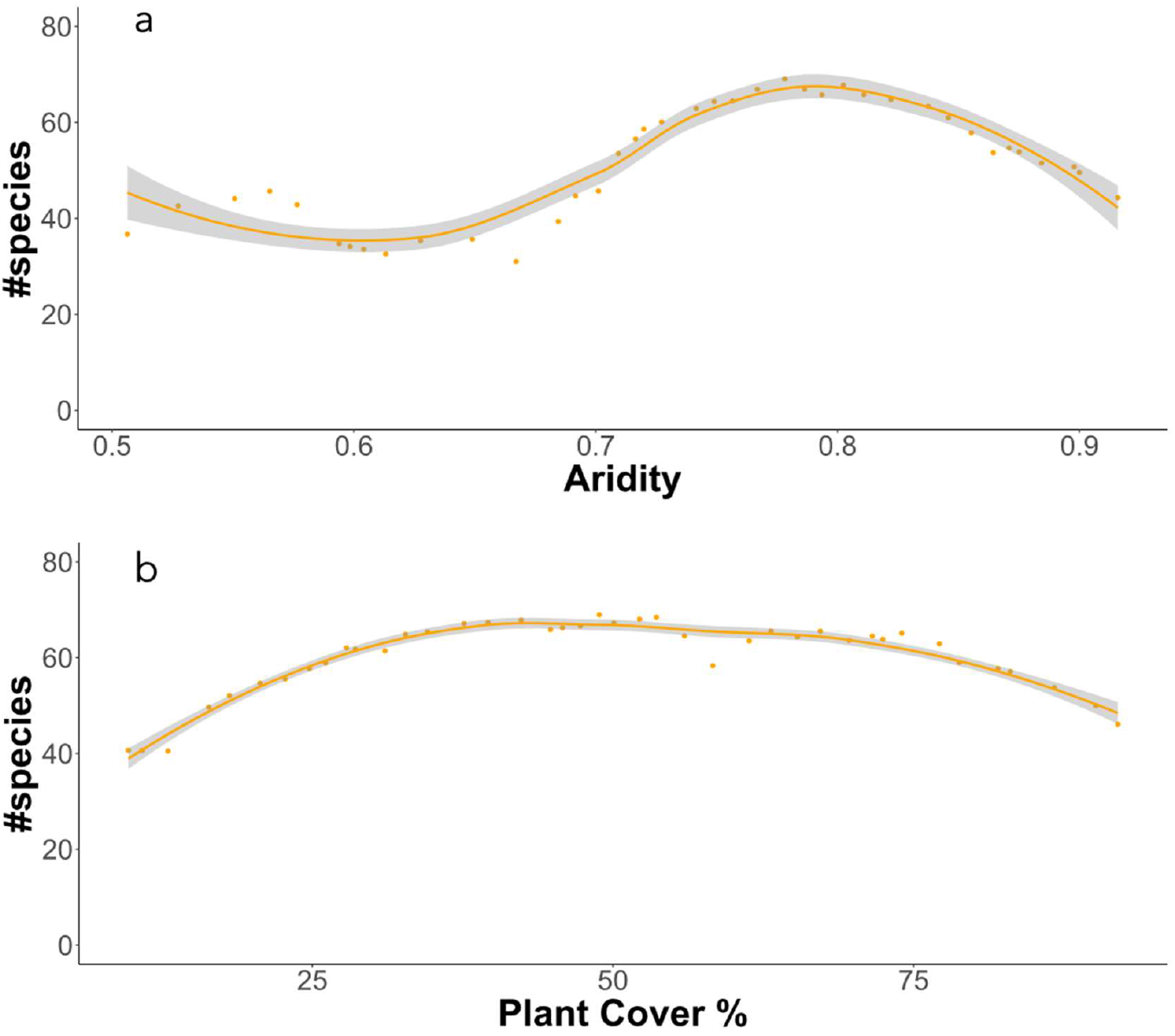
Average number of species observed in each aridity (**a**) and plant cover intervals (**b**) in Gross et al. 2024.

